# Modeling mechanisms of tremor reduction for essential tremor using symmetric biphasic DBS

**DOI:** 10.1101/585117

**Authors:** Shane Lee, Wael F Asaad, Stephanie R Jones

## Abstract

Essential tremor (ET) is the most common movement disorder, in which the primary symptom is a prominent, involuntary 4–10 Hz movement. For severe, medication refractory cases, deep brain stimulation (DBS) targeting the ventral intermediate nucleus of the thalamus (VIM) can be an effective treatment for cessation of tremor and is thought to work in part by disrupting tremor frequency oscillations (TFOs) in VIM. However, DBS is not universally effective and may be further disrupting cerebellar-mediated activity in the VIM. Here, we applied biophysically detailed computational modeling to investigate whether the efficacy of DBS is affected by the mechanism of generation of TFOs or by the pattern of stimulation. We simulated the effects of DBS using standard, asymmetric pulses as well as biphasic, symmetric pulses to understand biophysical mechanisms of how DBS disrupts TFOs generated either extrinsically or intrinsically. The model results suggested that the efficacy of DBS in the VIM is affected by the mechanism of generation of TFOs. Symmetric biphasic DBS reduced TFOs more than standard DBS in both networks, and these effects were stronger in the intrinsic network. For intrinsic tremor frequency activity, symmetric biphasic DBS was more effective at reducing TFOs. Simulated non-tremor signals were also transmitted during symmetric biphasic DBS, suggesting that this type of DBS may help to reduce side effects caused by disruption of the cerebellothalamocortical pathway. Biophysical details in the model provided a mechanistic interpretation of the cellular and network dynamics contributing to these effects that can be empirically tested in future studies.

**Significance Statement:** Essential tremor (ET) is a common movement disorder, whose primary symptom is an involuntary rhythmic movement of the limbs or head. An area of the human thalamus demonstrates electrical activity that oscillates at the frequencies of tremor, and deep brain stimulation (DBS) in this area can reduce tremor. It is not fully understood how DBS affects tremor frequency activity in the thalamus, and studying different patterns of DBS stimulation may help to clarify these mechanisms. We created a computational model of different shapes of DBS and studied how they reduce different hypothesized generators of tremor frequency activity. A greater understanding of how DBS affects the thalamus may lead to improved treatments to reduce tremor and alleviate side effects in patients with ET.

## 1 Introduction

Essential Tremor (ET) is a heterogeneous neurological disorder primarily noted by a low frequency (4–10 Hz), involuntary “intention” tremor of the limbs [11,28,29]. Both motor and cognitive deficits have been observed in ET [25, 30, 45], and though deficits in the cerebellum have been implicated, the pathophysiology is multimodal and not fully understood [10–12,19,30,41,47]. For debilitating, medication-refractory cases of ET, high frequency deep brain stimulation (DBS) of the ventral intermediate nucleus of the thalamus (VIM)—the cerebellar motor area of the thalamus—is a therapy that can be successful in alleviating symptoms [6,37].

Standard, “monophasic” DBS is presently thought to work by stimulating axons at high frequencies, effectively blocking tremor frequency neural activity [33]. However, the mechanisms by which DBS alleviates tremor are not fully understood [2,10], and DBS can also induce side effects such as paresthesias, gait imbalance, and impaired motor adaptation [1,10,13]. Furthermore, if standard DBS acts to block tremor activity by entraining axonal output or afferent input to VIM, the high frequency stimulation may also be blocking non-tremor-related cerebellar information, resulting in some of the observed deficits related to cerebellar activity [10]. Novel stimulation protocols therefore strive to selectively reduce pathological tremor frequency neural activity, process non-tremor input activity, and reduce side effects. Recently, feasibility and safety studies have investigated the use of DBS pulse sequences in which both the active and recharge phases are amplitude-balanced to treat movement disorders [3,4]. However, the efficacy and mechanisms by which this biphasic DBS act are not fully known.

Prior computational modeling has investigated multiple aspects of the timing of DBS on thalamic cells, including rebound bursting [8,9,26,48], and recent modeling has been focused on understanding tremor frequency oscillations (TFOs) in the VIM and their relation to DBS [27,51]. We recently developed a detailed computational model of a tremor frequency generating network within the VIM to show how network biophysical properties affected tremor and non-tremor activity [27]. In that work, the model simulated two distinct mechanisms of generation of TFOs in VIM: extrinsically-driven and intrinsically-generated. Extrinsic TFOs were driven by an external, non-VIM synaptic source such as the dentate nucleus of the cerebellum, while intrinsic TFOs were generated by the internal dynamics of VIM and thalamic reticular nucleus (TRN). Though some evidence exists for extrinsic TFOs, the role for intrinsic TFOs is not fully understood but may be crucially important in understanding the dynamics of both tremor as well as how DBS affects tremor. The prior modeling highlighted distinct network mechanisms involved in extrinsically and intrinsically generated TFOs and gave rise to experimentally testable predictions on how these oscillations may be modulated, particularly by pharmacology, to help improve future treatment strategies for ET.

Here, we sought to understand whether TFOs generated by distinct mechanisms responded differently to DBS and whether symmetric DBS simulated with cathode-anode biphasic pulse sequences may be more effective at reducing tremor or stimulation-induced side effects. We further establish a biophysical mechanism involving T-type Ca^2+^ by which biphasic stimulation reduces TFOs. Finally, we simulated a brief, non-tremor cerebellar signal during ongoing DBS to investigate whether transmission could be improved by different DBS pulse shapes while simultaneously reducing tremor-related activity.

## 2 Methods

We employed a previously developed computational model of the ventral intermediate nucleus of the thalamus (VIM) coupled to the thalamic reticular nucleus (TRN) [27] and extended it by modeling monopolar extracellular stimulation as DBS. In brief, the original model of a small tremor cluster in VIM consisted of 25 multicompartmental thalamocortical VIM cells and 25 multicompartmental TRN cells (Fig. 1A–C). If tremor clusters are responsible for tremor [39], targeted disruption of these clusters may be beneficial to alleviate tremor without affecting other aspects of VIM function. VIM cells in the model included both dendritic and axonal compartments, and the spiking at the most distal axonal site was analyzed to understand how tremor frequency activity in VIM affected downstream areas. Two modes of generation of TFOs were considered: extrinsically driven and intrinsically generated. In the absence of external stimulation, the network simulations in both models produced similar TFOs (example in Fig. 1D; see [27] for comparison). Aside from tonic currents and Poisson-distributed inputs applied to individual cells, in some simulations, a fast, excitatory input was simulated to reflect an error signal or other cerebellar or corticothalamic non-tremor input [5,46].

**Figure 1:**
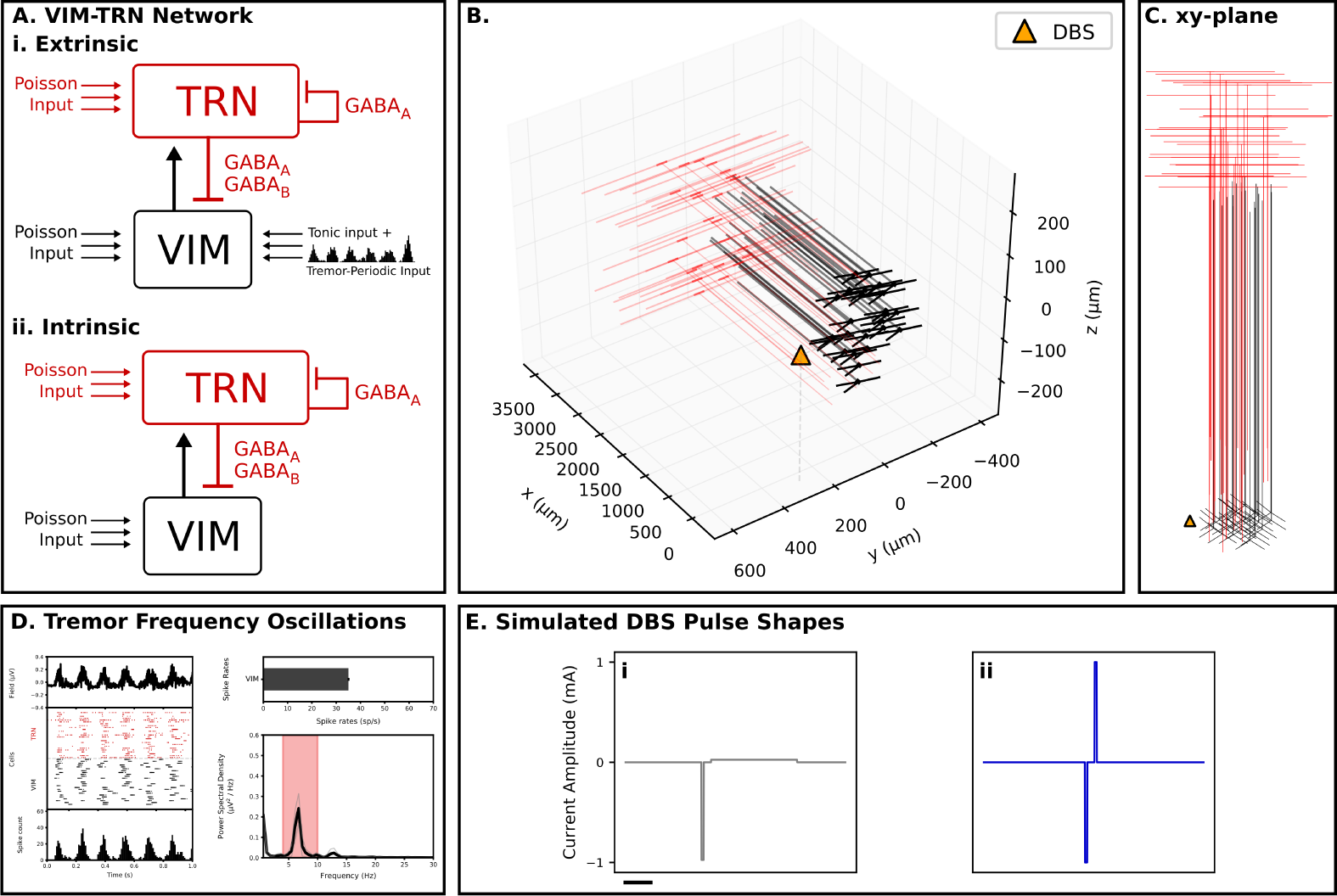
Model VIM-TRN networks with DBS. A. VIM-TRN Network schematic. 25 multicompartment VIM cells (black) and 25 multicompartment TRN cells (red) were simulated. Both VIM and TRN cells received Poisson distributed inputs. VIM-TRN connections were AMPA-ergic, while TRN-VIM connections were GABA_A_ and GABA_B_. TRN-TRN connections were GABA_A_. Ai. Extrinsic network received tonic applied current input as well as tremor-frequency-periodic AMPA-ergic synaptic input. Aii. Intrinsic network generated tremor-periodic oscillatory activity without external input. B. 3D perspective of network, with DBS stimulation location marked (orange triangle). C. xy-plane perspective of B. The center of mass of the VIM centered about the origin (0, 0, 0), with axons in the positive x direction toward the TRN. The “monopolar” extracellular stimulation, to simulate DBS, was placed at (250, 0, 0) *µm* (orange triangle). D. Without stimulation, both networks generated TFOs (example intrinsic shown, see [27]). Raster plot of VIM (black) and TRN (red) spikes for 1 s. VIM spike histogram at distal end of axon simulating output of VIM. Power spectrum for spike histogram shows TFOs. E. DBS pulse shapes simulated. Stimulation position, frequency, and primary pulse width were set for all simulations (see Methods). Current shapes and amplitudes were varied for different simulations. Ei. “Standard” DBS pulses consisted of a negative primary phase and a charge-balanced positive secondary phase of lower amplitude and longer width. Eii. DBS_*ca*_ consisted of a negative phase followed by a positive phase of equal amplitude. Scale bar for all is 1 ms.

### 2.1 DBS

Similar to prior studies [33], DBS was modeled as an extracellular monopolar stimulation through a homogeneous medium and best approximated a constant current stimulation. An extracellular potential was calculated based on the stimulation amplitude, transfer resistance through the medium, and distance between the stimulation and the compartment. The value of the extracellular resistivity was set to 350 Ω *cm* [24,33]. To simulate targeted stimulation to a tremor cluster, the simulated DBS electrode location was placed 250 *µm* along the y-axis, just off of the center of VIM (Fig. 1B–C), different from earlier work [33]. Transfer impedances were approximately 1 *k*Ω. Current amplitude at the stimulation site was specified in milliamps (mA).

Three different, charge balanced, monopolar DBS pulses were simulated (Fig. 1E and see Fig. 6A). For “standard” DBS, we simulated a cathodic active impulse, generating a negative current at the stimulation site, with a pulse width of 90 *µs* and a stimulation frequency of 130 Hz. These pulses were charge-balanced, with a balance pulse width that was calculated based on the entire cycle length at the stimulation frequency (approximately 7.69 ms).

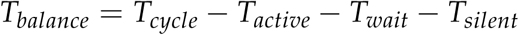

where *T*_*wait*_ was 300 *µs* and *T*_*silent*_ was 3.9 ms. The balance amplitude was therefore calculated such that:

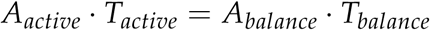

We also simulated charge-balanced, symmetric biphasic pulses, in which *A*_*active*_ equaled *A*_*balance*_ and *T*_*active*_ equaled *T*_*balance*_ [3,4,22]. These pulses were also simulated at 130 Hz, with pulse widths of both phases 90 *µs*. We simulated both cathodic-leading (DBS_*ca*_) and anodic-leading (DBS_*ac*_) (Fig. 1Eii and Fig. 6A).

### 2.2 Output of VIM model

The simulations were run similarly to our prior work [27]. Ten iterations were run for comparison across noise conditions, and sources of noise were identical to the original model and included the spatial locations of the VIM and TRN cells, initial membrane potentials, and realizations of the Poisson-distributed background synaptic inputs.

As in the previous model [27], the principal output was the axonal output spiking from the VIM cells. During DBS stimulation, the spike output from VIM cells was multimodal. To quantify this, we classified spiking as DBS-following (spike rates that exceeded 100 sp/s and often followed the DBS stimulation frequency), rhythmic spiking (spike rates that formed bursting-type cell responses, or low spiking (nearly zero spike rates in a specified interval time-locked to the onset of stimulation). Prior computational and experimental work has suggested that high frequency DBS leads to synaptic failure and reduces the probability of output for those cells [42]; therefore in these simulations, tremor frequency activity was calculated specifically from the non-high frequency spiking cells.

Spike counts of different spike types were compared for different DBS conditions. In these simulations, we varied the strength of DBS amplitude for different pulse patterns. We compared the output spike types at a given DBS amplitude compared to control with no DBS stimulation. To assess significance, we used Fisher’s Exact test for 2×2, 3×2, or 3×3 matrices (types of cells *x* conditions).

For different DBS types, we analyzed evoked spiking responses due to a single, fast, excitatory synaptic input of varying strength. DBS was turned on at 0.150 s, and the input was delivered at 0.4 s. These simulations were run for a total of 0.6 s of simulation time. The evoked response was defined as the quadrature of the convolved VIM spike output in the 50 ms following the stimulation input in cells that were non-high frequency spiking. A control condition was run in which the strength was set to 0, and the analysis was performed identically.

### 2.3 T-type Ca^2+^ events

The number of discrete T-type Ca^2+^-mediated current events was analyzed to quantify whether T-type Ca^2+^ mechanisms were employed during low spiking versus rhythmic cells. Events were counted by rectifying the T-type Ca^2+^ current, downsampling from 100 kHz to 1 kHz, and finding peaks that passed a current threshold of 5 × 10^*-*3^*µA*/*cm*^2^. This method captured substantial current events that occurred when both the *m* and *h* gates were co-active and was consistent with manual rating. More T-type Ca^2+^ current events suggested this current’s involvement in the spiking patterns observed. Distributions of T-type Ca^2+^-mediated current events were compared between rhythmic spiking cells versus low spiking cells for each DBS pulse pattern, at each different DBS amplitude. Comparisons were made only when there were 10 or greater cells in both categories.

### 2.4 Statistics

For most comparisons, a non-parametric Mann-Whitney U test was used to assess significance [32,50], which for all simulations was defined as *p* < 0.025. For simulations with multiple comparisons, we ran a Benjamini-Hochberg test [7] with a false positive rate (*q*) of 0.05. Most analyses were conducted using Python 3.5 with standard modules, SciPy 0.18, and NumPy 1.13. The Fisher’s exact test [14,34] was performed using R 3.4. All error is reported as standard error.

A step size of 0.010 ms, smaller than the prior work of 0.025 ms, was used for all simulations here to provide finer temporal resolution during the short active pulses due to DBS stimulation. The original model along with the DBS stimulation introduced here is available on ModelDB (https://senselab.med.yale.edu/ModelDB/) with accession number 240113.

## 3 Results

### 3.1 Mechanisms of tremor reduction differ for extrinsic and intrinsic networks

Effects of DBS pulse shape and amplitude were modeled on two mechanisms of TFOs, extrinsic and intrinsic (Fig. 1). TFOs in both networks were significantly reduced by all simulated DBS types, but individual cells in each network exhibited differing responses.

#### 3.1.1 Extrinsic Network

In the extrinsic network, both DBS types significantly lowered TFOs compared to a no-stimulation baseline, from 0.5–2.0 mA (Table 1A and Fig. 2A, significance denoted by gray + for standard DBS, blue + for DBS_*ca*_). DBS_*ca*_ also significantly lowered TFOs at 0.25 mA, suggesting that biphasic may be effective at low amplitudes. In contrast, at 0.25 mA, standard DBS actually increased TFOs, demonstrating that extrinsically generated TFOs can paradoxically increase with DBS stimulation. This occurred because the low amplitude DBS activity was not enough to overcome the inactivation of T-type Ca^2+^ necessary to reduce TFOs but rather contributed to amplifying VIM cell bursts. The lack of entrainment and persistent bursting of cells is seen in the example rasters in Fig. 2Aii.

**Table 1:**
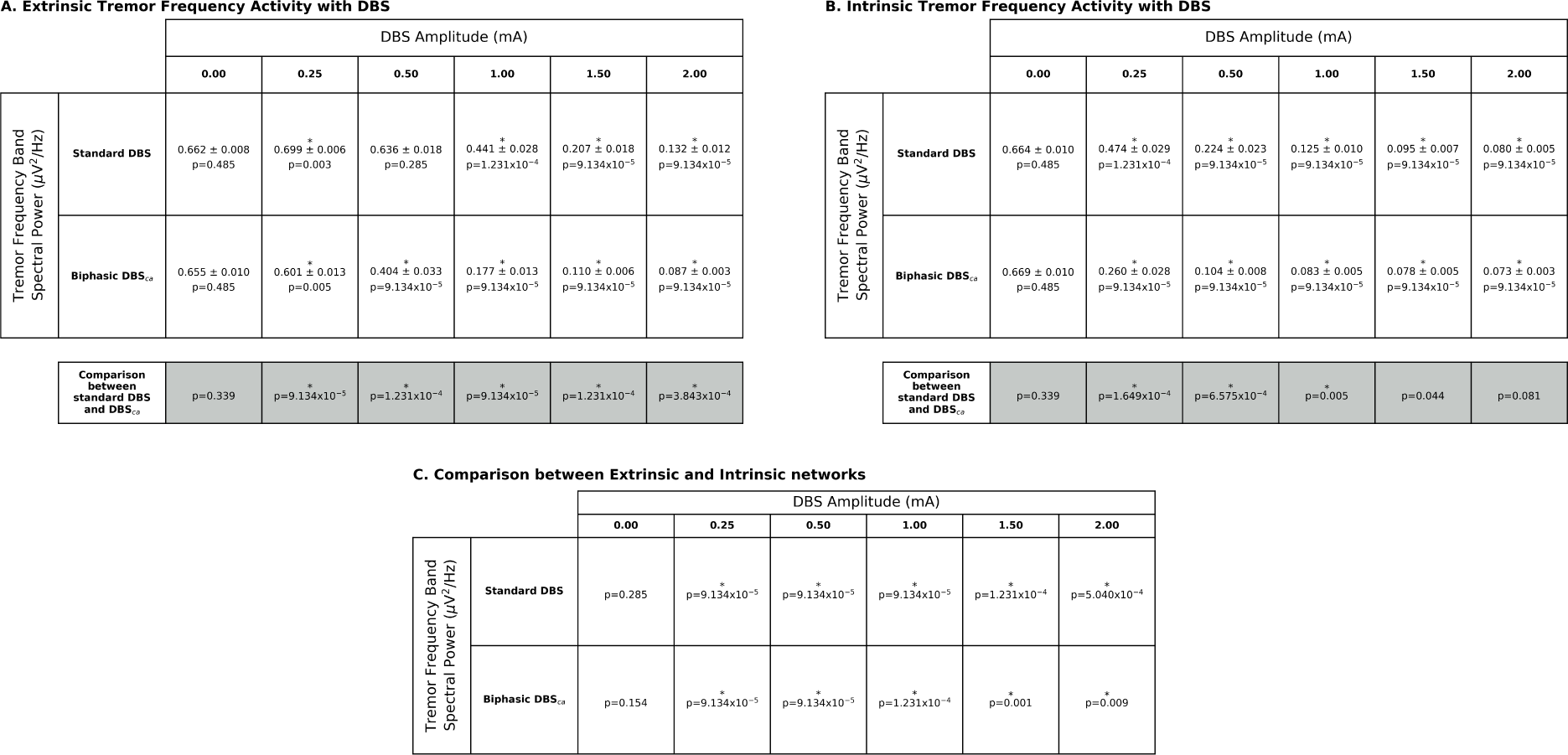
Tremor frequency power for different networks and DBS types. Tremor frequency power was measured for extrinsic (A) and intrinsic (B) during standard DBS and DBS_*ca*_. Tested amplitudes (columns) for a given stimulation pattern (rows). Top rows (white) show mean spectral power for TFOs (n=10, standard error reported). Comparisons within a row are to the baseline at 0.00 mA. Last rows (gray) are same-amplitude comparisons of tremor frequency power between standard DBS and DBS_*ca*_. C. Comparison between extrinsic and intrinsic networks for each amplitude and stimulation type. See Fig. 2.

**Figure 2:**
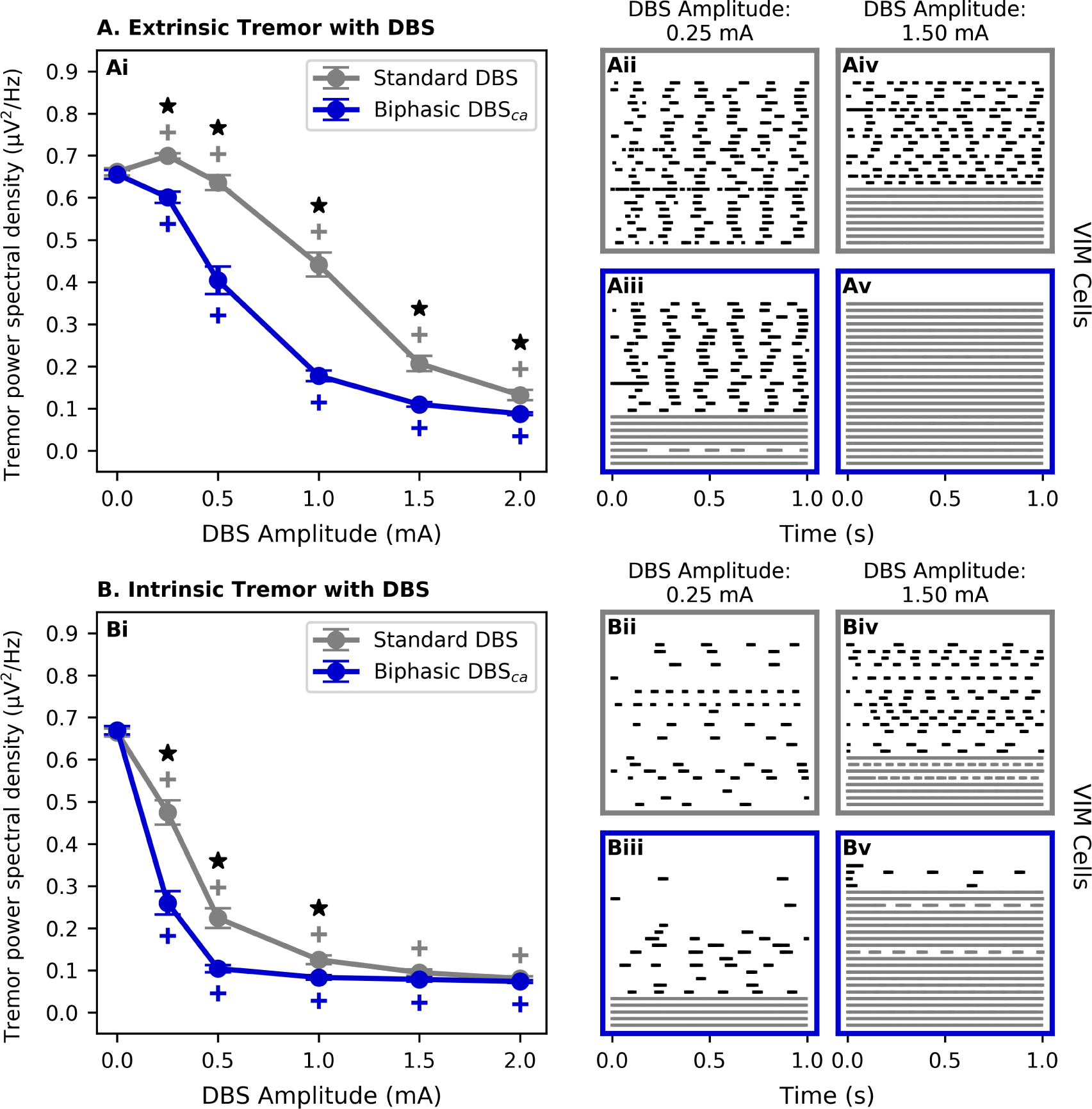
Model DBS reduced intrinsic TFOs more than extrinsic; biphasic DBS reduced TFOs in both models at lower amplitudes compared to standard. Effects of DBS were simulated for both extrinsic (A) and intrinsic (B) networks. Gray + denotes significantly different power for standard DBS compared to no DBS at that amplitude. Blue + denotes significantly different power for DBS_*ca*_ DBS compared to no DBS at that amplitude. Black star denotes significantly different power between standard DBS and DBS_*ca*_. A. DBS with extrinsic TFOs. Increased amplitude of standard DBS (gray) and DBS_*ca*_ (blue) reduced tremor oscillations significantly compared to baseline. DBS_*ca*_ reduced TFOs less than standard DBS for all non-zero amplitudes of stimulation. Aii–Aiii. DBS_*ca*_ induced more high frequency spiking cells than standard DBS at 0.25 mA. Example spike rasters shown. Aiv–Av. DBS_*ca*_ induced high frequency spiking in all cells at 1.5 mA, while standard DBS resulted in both high frequency spiking and uncorrelated rhythmic bursting. B. DBS with intrinsic TFOs. Increased amplitude of DBS reduced TFOs significantly compared to baseline for both stimulation patterns. DBS_*ca*_ reduced TFOs significantly greater than standard DBS at 0.25, 0.5, and 1.0 mA. Bii–Biii. Example spike rasters for each DBS pattern, shown for 0.25 mA, where no cells were high frequency with standard DBS while DBS_*ca*_ induced some. Biv–Bv. Example spike rasters at 1.5 mA. High frequency spiking was observed in some cells with standard DBS, but high frequency spiking was induced in the majority of cells by DBS_*ca*_. See Fig. 3 and Table 2.

In a direct comparison with standard DBS, biphasic DBS resulted in significantly lower TFOs at all tested amplitudes (Fig. 2A, significance denoted by black stars), further suggesting that biphasic pulse shapes may be more effective at lowering tremor (gray boxes in Table 1A).

**Table 2:**
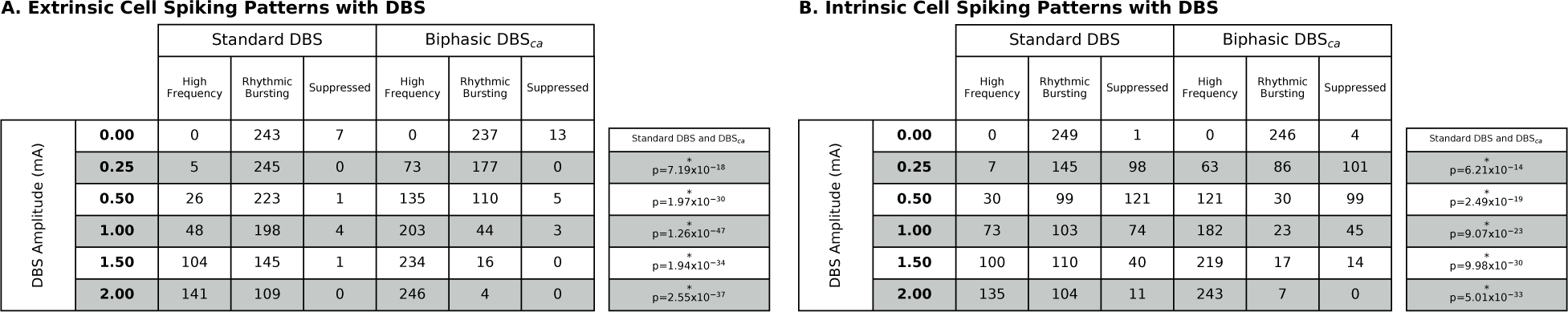
DBS-induced VIM cell spiking patterns for both extrinsic and intrinsic networks. For each network, each DBS type (Standard and DBS_*ca*_) was simulated. Cells were classified by spike patterns: high frequency, rhythmic bursting, and suppressed cells (see Methods). Twenty-five VIM cells were tested during 10 simulations; the totals in each row equal 250 for each DBS stimulation. Fisher’s Exact tests used to assess significance. See Fig. 3. A. Among non-high frequency cells, the extrinsic network generated overwhelmingly rhythmic bursting cells. Compared to standard DBS, DBS_*ca*_ led to significantly more high frequency spiking cells. B. Among non-high frequency cells, the intrinsic network generated both rhythmic bursting and suppressed spiking cells. Compared to standard DBS, DBS_*ca*_ led to significantly more high frequency cells.

Example rasters for the axonal spiking output of the VIM cells demonstrated how different cells responded to DBS to result in lowered TFOs in the extrinsic network (Fig. 2Aiii–Av). At low DBS amplitudes, biphasic DBS entrained high frequency spiking in a small population of cells (Fig. 2Aiii), consistent with previous explanations of tremor frequency interference. However, the remaining cells had rhythmic and nearly synchronous burst spiking, similar to the network during standard DBS. At 1.5 mA, standard DBS led to both a population of high frequency spiking cells but also a majority of rhythmic but less synchronous, decorrelated cells (Fig. 2Aiv), while DBS_*ca*_ entrained high frequency spiking in nearly all VIM cells (Fig. 2Av).

#### 3.1.2 Intrinsic Network

DBS affected the intrinsic network differently than the extrinsic network. TFOs were significantly reduced at all tested amplitudes, for both standard DBS and DBS_*ca*_(Table 1B and Fig. 2B, significance denoted by gray + for standard DBS, blue + for DBS_*ca*_). The reduction was significantly greater for DBS_*ca*_ compared to standard DBS at 0.25, 0.5, and 1.0 mA, further suggesting that biphasic stimulation may be more efficient in reducing tremor (black stars in Fig. 2B and gray boxes in Table 1B).

At low stimulation (0.25 mA), standard DBS reduced TFOs in the intrinsic network by completely suppressing spiking in some cells and decorrelating burst spiking in the remaining cells (Fig. 2Bii). DBS_*ca*_ also entrained the spiking of multiple cells, even at the lowest tested amplitude (Fig. 2Biii). At 1.5 mA, high frequency spiking is seen in a subset of cells excited by standard DBS, with the remaining cells mostly bursting in an uncorrelated manner. In contrast, DBS_*ca*_ at 1.5 mA led to high frequency spiking in the majority of cells, while the remaining cells were mostly silent (Fig. 2Bv).

For all tested amplitudes of stimulation for both standard DBS and DBS_*ca*_, reductions in TFOs were significantly greater in the intrinsic network compared to the extrinsic network. Though it is not fully known to what extent the intrinsic oscillatory activity of VIM contributes to the expression of tremor [27], these results suggest that DBS reduces intrinsic TFOs more than extrinsic and by distinct mechanisms. The distinguishable spike outputs provide testable predictions on the mechanisms generating TFOs, when perturbed by different DBS stimulation pulses. Furthermore, for both networks there were a range of stimulation amplitudes in which DBS_*ca*_ reduced TFOs more than standard DBS.

### 3.2 Distinct VIM cell responses to DBS in extrinsic and intrinsic networks

The spike rasters from example cells in Fig. 2 showed that compositions of cells with different spike patterns—rhythmic bursting, high frequency spiking, and silent— differed depending on the network and DBS protocol. We quantified whether the composition of cells was systematically different due to the type of DBS in both extrinsic and intrinsic networks (Fig. 3 and Table 2).

**Figure 3:**
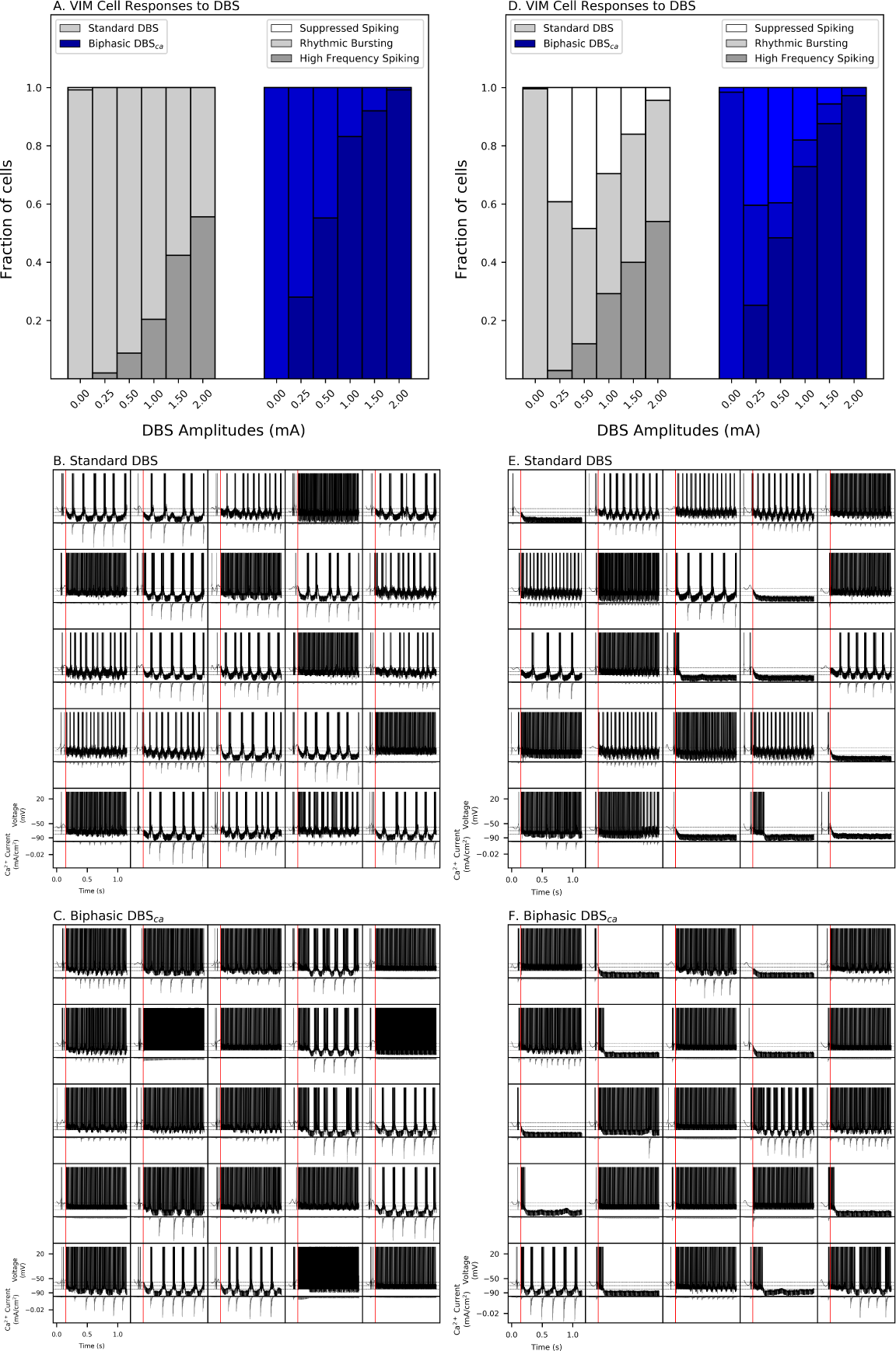
DBS creates different patterns in different networks. Both standard DBS and DBS_*ca*_ gave rise to rhythmic bursting in both the extrinsic and intrinsic networks. Suppressed spiking was also observed during intrinsic but extrinsic TFOs. Rhythmic spiking was always associated with increased T-type Ca^2+^ current amplitude. Examples of spiking explored further in Fig. 4. A–C. Extrinsic network with DBS. A. Summary of VIM cell responses to different amplitudes of standard DBS (gray) and DBS_*ca*_ (blue). Cells were sorted by spike rate: high frequency spiking (dark shade), rhythmic bursting (medium shade), and suppressed spiking (light shade, not observed here). For all DBS types, as DBS amplitude was increased, the fraction of DBS cells increased. B–C. Examples of all VIM cells of a single trial during standard DBS (B) and DBS_*ca*_ (C), at 1.0 *mA*. DBS was initiated at the red line simultaneously for all cells. Rhythmic bursting always involved T-type Ca^2+^ currents, which is shown for each VIM cell soma beneath the voltage plot. D–F. Intrinsic network with DBS. D. DBS_*ca*_ resulted in greater numbers of high frequency spiking cells compared to standard DBS. Standard DBS gave rise to more rhythmic bursting, while both biphasic sequences resulted in fewer rhythmic bursting cells. Rhythmic bursting was always associated with increased T-type Ca^2+^ current amplitude. Both DBS types also resulted in suppressed spiking cells not seen during DBS in the extrinsic network. E–F. Examples of all VIM cells of a single trial during standard DBS (B) and DBS_*ca*_ (C) at 1.0 *mA*. Rhythmic bursting always involved T-type Ca^2+^ currents.

For extrinsically-driven TFOs, all three DBS types led to high frequency spiking output in the VIM cells as well as decorrelated rhythmic bursting in other cells (Fig. 3A–C and Table 2A). Almost no suppressed cells were observed in these networks with DBS. Example output for 25 VIM cells in a single network is shown during standard DBS (Fig. 3B) and DBS_*ca*_ (Fig. 3C), all demonstrating similar phenomena. For all tested non-zero DBS amplitudes, both biphasic DBS types led to significantly greater fractions of high frequency spiking cells than standard DBS (Table 2A).

For intrinsic networks, DBS reduced intrinsically-generated TFOs with high frequency spiking but also suppressed spiking in some cells (Fig. 3D and Table 2B), which was not seen in the extrinsic network. While standard DBS led to a mixture of sup-pressed and rhythmic bursting cells (Fig. 3E), DBS_*ca*_ generally suppressed spiking activity and did not generate as much rhythmic bursting (Fig. 3F). DBS_*ca*_ led to significantly more high frequency VIM cells compared to standard DBS (Table 2B).

The mechanisms by which DBS reduced TFOs in the model VIM was therefore dependent upon the mechanism generating the TFOs. While DBS reduced TFOs in extrinsic networks, it did so principally by increasing the number of high frequency—often DBS-entrained—spiking, which was more pronounced with DBS_*ca*_ than standard DBS. In contrast, suppression of spiking was observed in intrinsic networks with both DBS types. While standard DBS also led to rhythmic bursting, only DBS_*ca*_ simultaneously reduced TFOs and suppressed spiking output of non-high-frequency spiking cells.

### 3.3 Role of T-type Ca^2+^ currents in suppression of VIM cell spiking in intrinsic TFOs

We further investigated the suppression of VIM cells during DBS on intrinsic TFOs and found that it was related to T-type Ca^2+^ inactivation. Somatic T-type Ca^2+^ currents were observed for rhythmic bursting in the network (Fig. 3D–F, gray traces below voltages), while these currents were not present during suppressed responses. For both stimulation patterns (Standard and DBS_*ca*_) at each DBS amplitude tested (0.00– 2.00 mA), the number T-type Ca^2+^ current events were counted (see Methods). Higher numbers of T-type Ca^2+^ events suggested greater involvement in generating the activity, as the events were time-locked to the spiking activity. For all valid comparisons at a given pattern at a specific amplitude, rhythmic bursting cells were associated with significantly greater T-type Ca^2+^ current events than the suppressed cells (Table 3). In nearly all cases, suppressed cells had near-zero T-type Ca^2+^ events.

**Table 3:**
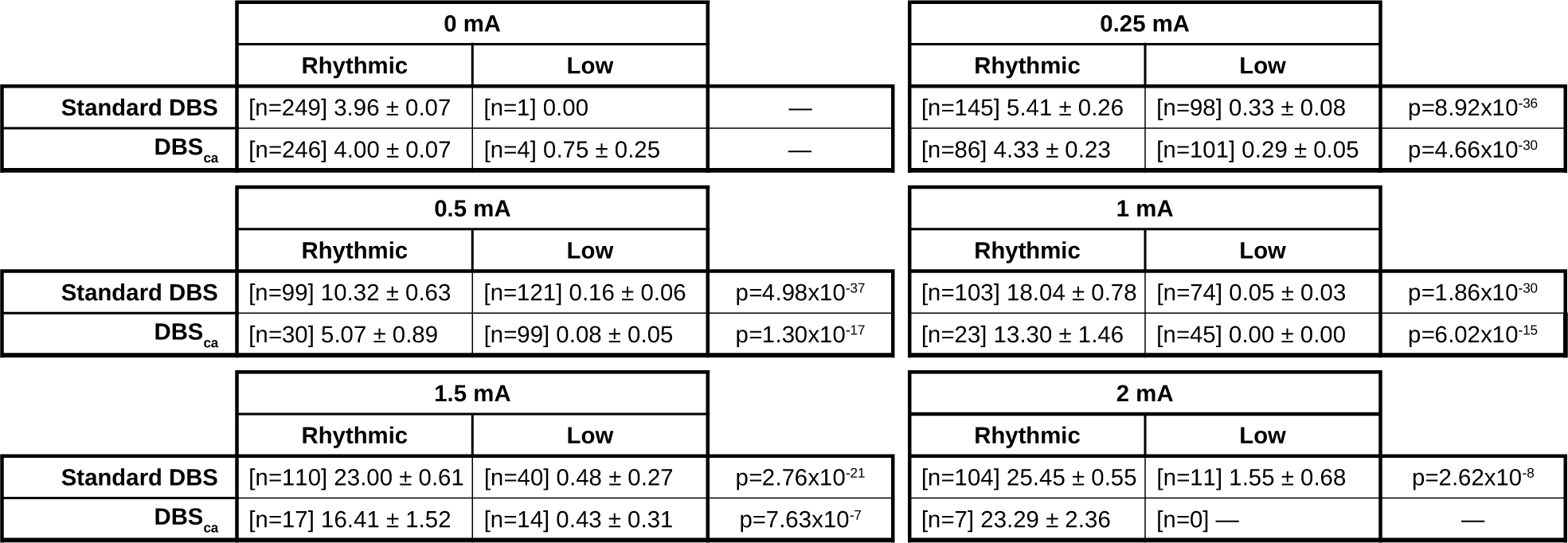
T-type Ca^2+^ current events during DBS for non-high frequency cells. T-type Ca^2+^ events were counted for non-high frequency cells and separated into rhythmic bursting and suppressed spiking types (see Methods). For all simulations in which there were at least 10 rhythmic bursting and sup-pressed spiking types, rhythmic bursting cells had significantly greater T-type Ca^2+^ current events than the suppressed spiking cells. See Fig. 3.

|  | 0 mA |  |  |
| --- | --- | --- | --- |
|  | Rhythmic | Low |  |
| Standard DBS | [n=249] $3.96 \pm 0.07$ | [n=1] 0.00 | — |
| DBS<sub>ca</sub> | [n=246] $4.00 \pm 0.07$ | [n=4] $0.75 \pm 0.25$ | — |
|  | 0.5 mA |  |  |
|  | Rhythmic | Low |  |
| Standard DBS | [n=99] $10.32 \pm 0.63$ | [n=121] $0.16 \pm 0.06$ | $p=4.98 \times 10^{-37}$ |
| DBS<sub>ca</sub> | [n=30] $5.07 \pm 0.89$ | [n=99] $0.08 \pm 0.05$ | $p=1.30 \times 10^{-17}$ |
|  | 1.5 mA |  |  |
|  | Rhythmic | Low |  |
| Standard DBS | [n=110] $23.00 \pm 0.61$ | [n=40] $0.48 \pm 0.27$ | $p=2.76 \times 10^{-21}$ |
| DBS<sub>ca</sub> | [n=17] $16.41 \pm 1.52$ | [n=14] $0.43 \pm 0.31$ | $p=7.63 \times 10^{-7}$ |
|  | 0.25 mA |  |  |
|  | Rhythmic | Low |  |
| Standard DBS | [n=145] $5.41 \pm 0.26$ | [n=98] $0.33 \pm 0.08$ | $p=8.92 \times 10^{-36}$ |
| DBS<sub>ca</sub> | [n=86] $4.33 \pm 0.23$ | [n=101] $0.29 \pm 0.05$ | $p=4.66 \times 10^{-30}$ |
|  | 1 mA |  |  |
|  | Rhythmic | Low |  |
| Standard DBS | [n=103] $18.04 \pm 0.78$ | [n=74] $0.05 \pm 0.03$ | $p=1.86 \times 10^{-30}$ |
| DBS<sub>ca</sub> | [n=23] $13.30 \pm 1.46$ | [n=45] $0.00 \pm 0.00$ | $p=6.02 \times 10^{-15}$ |
|  | 2 mA |  |  |
|  | Rhythmic | Low |  |
| Standard DBS | [n=104] $25.45 \pm 0.55$ | [n=11] $1.55 \pm 0.68$ | $p=2.62 \times 10^{-8}$ |
| DBS<sub>ca</sub> | [n=7] $23.29 \pm 2.36$ | [n=0] — | — |

Example differences in T-type Ca^2+^ current for the VIM cell spike types under DBS are shown in Fig. 4. Rhythmic cells demonstrated strong T-type Ca^2+^ currents that led to bursting behavior in the soma and output (Fig. 4A). In contrast, depolarization was sustained in high frequency cells (Fig. 4B). Though sometimes these cells had bouts of T-type Ca^2+^ activity, it was not necessary to sustain high frequency activity. Even when the soma did not elicit action potentials, the depolarization was enough to elicit spiking in the output at the axon terminals. Suppressed cells exhibited no T-type Ca^2+^ activity, as the T-type Ca^2+^ channels were inactivated (Fig. 4C). Subthreshold somatic activity driven by the DBS stimulation did not elicit spiking in either the soma or the axonal outputs.

**Figure 4:**
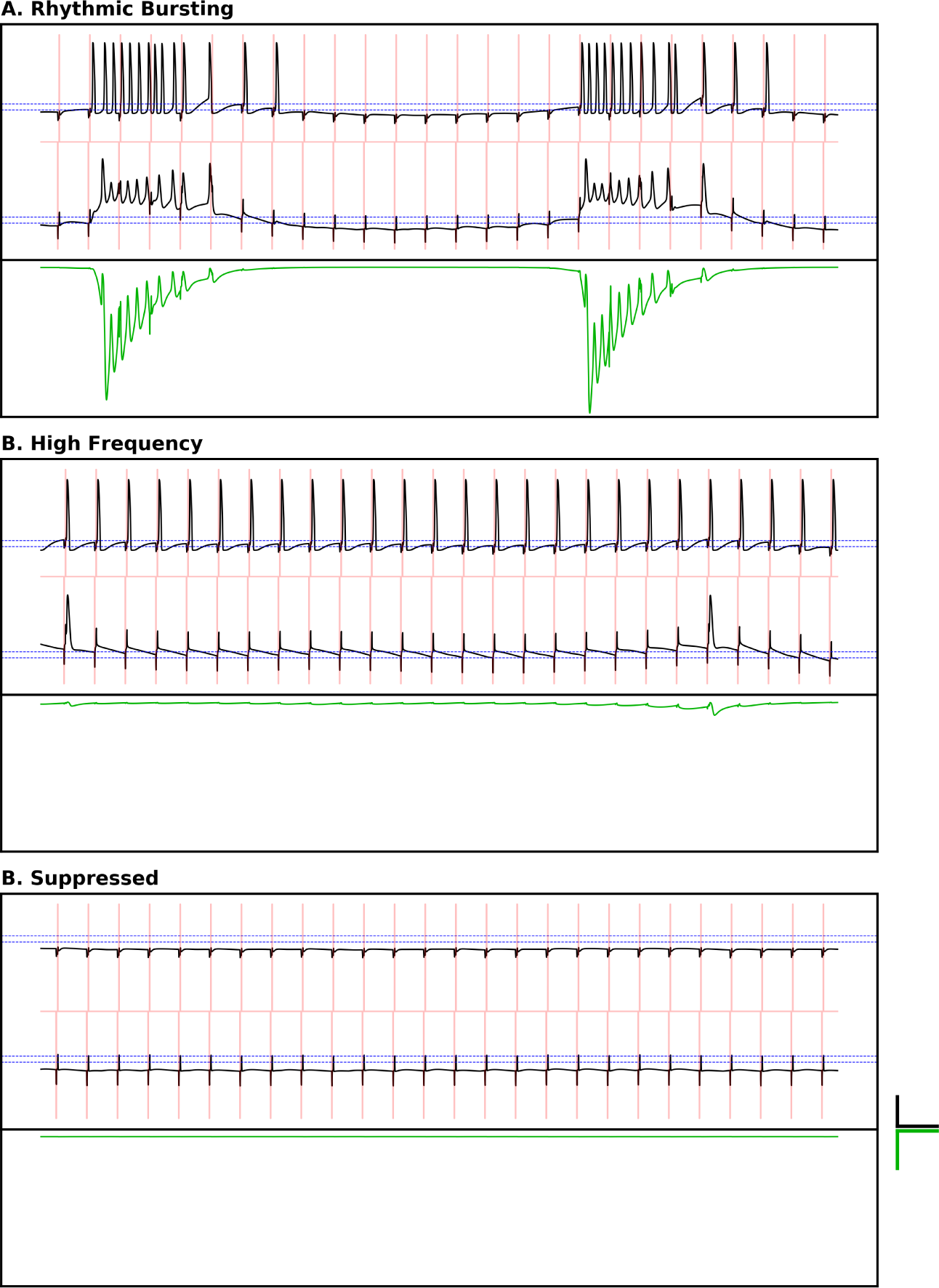
Spiking patterns observed with DBS. Three main spiking patterns were observed with DBS. Top panels are voltage traces of 2 compartments: axonal compartment on top, somatic below. Red trace is the DBS_*ca*_ stimulation. Similar patterns were also observed during standard DBS. Blue lines denote −74 mV and −64 mV. Lower panels show T-type Ca^2+^ current (green). Voltage scale is 50 mV. Current scale is 5 × 10^*-*3^ *µA*/*cm*^2^. Temporal scale is 10 ms. DBS in the extrinsic networks consisted of both high frequency and rhythmic responses; intrinsic networks also demonstrated suppressed responses. A. Rhythmic spiking consisted of de-inactivated T-type Ca^2+^ at the soma that transmitted to spiking at the axon. High T-type Ca^2+^ current is associated with the bursts. B. High frequency spiking (often DBS-entrained) occurred for high amplitudes of stimulation but also when the somatic stimulation was always in a voltage range above the inactivation threshold of T-type Ca^2+^. Little T-type Ca^2+^ current is activated here. C. Suppression occurred when the somatic membrane potential was inactivated. No T-type Ca^2+^ current is activated here.

These data suggested that DBS led to suppressed cells by maintaining a steady hyperpolarization through continued inactivation of T-type Ca^2+^ currents, which may be a therapeutic strategy to suppress pathological TFOs rather than through evoking high frequency spiking activity. Among non-high frequency spiking cells, standard DBS led to both suppressed spiking and rhythmic bursting, while DBS_*ca*_ led primarily to sup-pressed spiking, suggesting that the biphasic pulses may have helped to increase the probability of the cell membrane potential in the inactivated state that silenced T-type Ca^2+^ activity.

### 3.4 Biphasic DBS enabled inputs to be transmitted through the intrinsic network

When biphasic DBS was applied to the intrinsic model network, a substantial fraction of the non-DBS-following VIM cells were suppressed (Fig. 3D–E). Because the cerebellum is involved in motor control and error processing [10], we tested whether these suppressed cells could serve as a template to help enable salient, non-tremor inputs be transmitted through the VIM toward cortical targets.

We simulated a non-tremor, fast, excitatory input to both the extrinsic and intrinsic networks during DBS. Evoked spike rates for 50 ms following the input were measured in VIM cells. Cells with high frequency spiking induced by DBS were excluded from this analysis. For both the extrinsic and intrinsic networks, non-tremor, external inputs evoked response spike rates were measured in non-high-frequency cells in the 50 ms following the input during both ongoing DBS types, each at 1.0 mA, with two different strengths of input compared to no input: 0.02 *µS*, and 0.04 *µS* (Fig. 5, see Methods).

**Figure 5:**
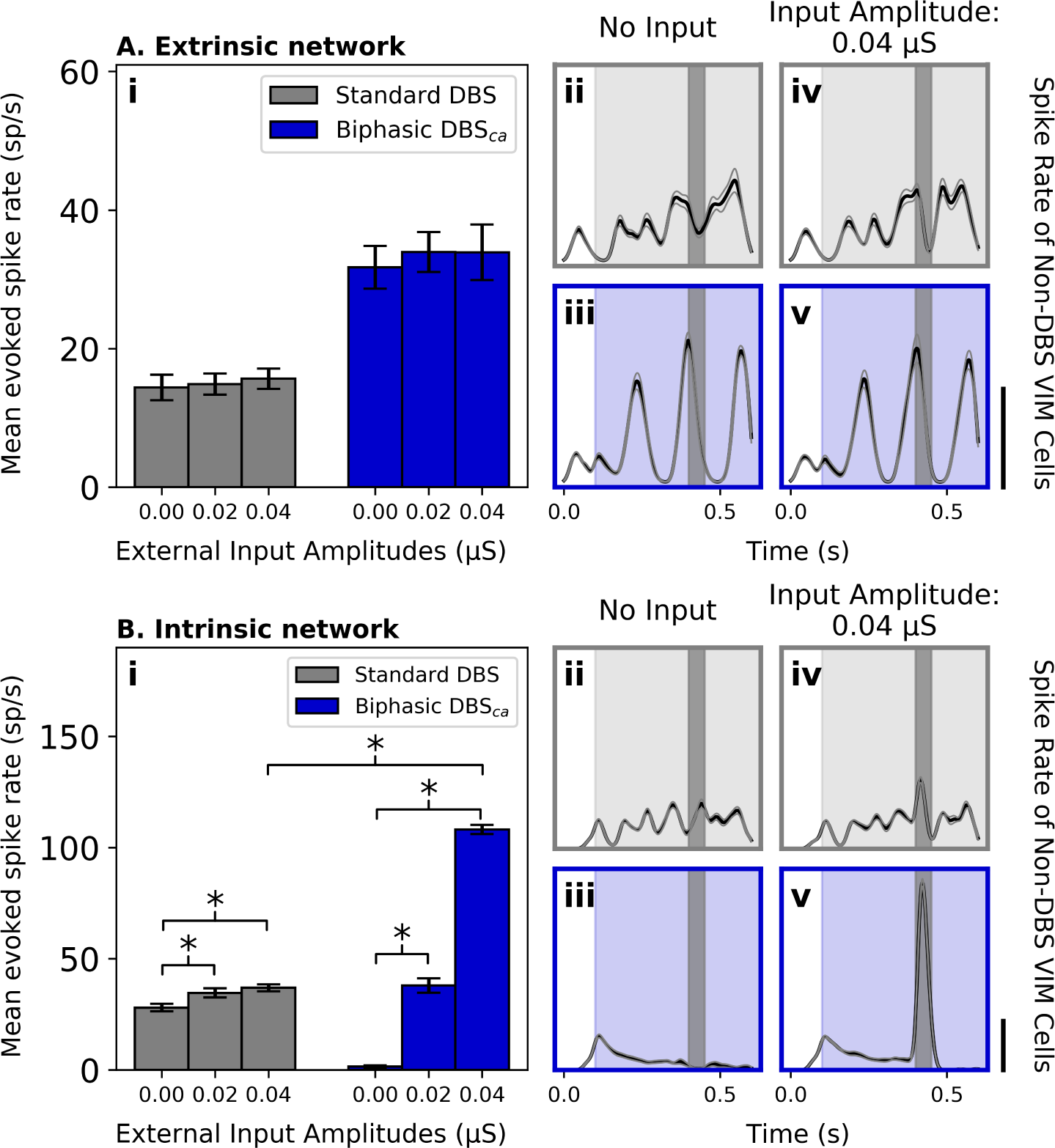
External synaptic inputs significantly increased the output rate during DBS, only for the intrinsic network. Rapid input from an area external to VIM during different mechanisms of TFOs with different DBS pulse shapes was simulated with a brief AMPA-ergic synaptic input was simulated to the VIM cells. External input appeared at 0.4 s. Evoked interval shaded in dark gray. Standard DBS (gray) and DBS_*ca*_ (blue) were tested. Scale bars for evoked rates are 40 sp/s. A. Extrinsic networks during DBS did not increase spike rate significantly due to external synaptic input, for either input strength. Ai. Evoked spike rates during each DBS type due to the external synaptic input. No significant increase resulted from either strength of input or DBS type. Aii–Aiii. Baseline aggregate spike rates of VIM cells with no input. DBS stimulation on during shaded interval. Aiv–Av. Aggregate spike rates of VIM cells with an input amplitude of 0.04 *µS* showed no significant increases following evoked input. B. Intrinsic networks during DBS showed significant increases in evoked spike rates due to external synaptic input. Bi. Evoked spike rates during both DBS types due to external input. During both DBS types, both tested inputs resulted in significant increases in evoked rates. DBS_*ca*_ resulted in significantly higher evoked rates at 0.04 *µS* compared to standard DBS. Bii–Biii. Baseline aggregate spike rates of VIM cells with no input. Biv–Bv. Input amplitude of 0.04 *µS* showed a small but significant increase during standard DBS but a large increase during DBS_*ca*_.

None of the external inputs evoked a significant increase in VIM cell spike rates during DBS for the extrinsic network (Fig. 5A and Table 4A), indicating that external inputs would not be propagated in networks with extrinsically driven TFOs. In contrast, external inputs evoked significant responses in VIM cells during all DBS types in the intrinsic network (Fig. 5B and Table 4B). Within DBS types, each input evoked significant responses compared to baselines.

**Table 4:** Evoked responses during different DBS types for both networks. Evoked responses in the 50 ms following strong “cerebellar” synaptic inputs were measured during ongoing DBS. Also see Fig. 5. In first 2 rows, evoked rates (sp/s) within stimulation types were compared to baseline at 0 *µS*. Last row compares evoked responses between standard DBS and DBS_*ca*_. A. Extrinsic network. As input amplitude increased, no significant changes were seen in the evoked response compared to baseline, for any of the DBS types. B. Intrinsic network. Inputs significantly increased evoked spike rates for all DBS types. DBS_*ca*_ had a significantly lower baseline response compared to standard DBS and a significantly higher evoked response at 0.04 *µS*.

**A. Extrinsic Evoked Responses during DBS**
| Evoked Responses (sp/s) for<br>Different DBS Pulse Shapes | | Input Amplitudes (μS) | | |
| --- | --- | --- | --- | --- |
|  |  | 0.00 | 0.02 | 0.04 |
| | Standard DBS | 14.373 ± 1.838 | 14.865 ± 1.542<br>p=0.339 | 15.648 ± 1.478<br>p=0.312 |
| | DBS<sub>ca</sub> | 31.733 ± 3.078 | 33.925 ± 2.892<br>p=0.396 | 33.890 ± 4.006<br>p=0.339 |
| Comparison between standard DBS and DBS<sub>ca</sub> | | p=5.040$\times 10^{-4}$* | p=1.231$\times 10^{-4}$* | p=0.001* |

**B. Intrinsic Evoked Responses during DBS**
| Evoked Responses (sp/s) for<br>Different DBS Pulse Shapes | | Input Amplitudes (μS) | | |
| --- | --- | --- | --- | --- |
|  |  | 0.00 | 0.02 | 0.04 |
| | Standard DBS | 27.986 ± 1.691 | 34.627 * ± 2.007<br>p=0.019 | 36.912 * ± 1.559<br>p=3.843$\times 10^{-4}$* |
| | DBS<sub>ca</sub> | 1.507 ± 0.497 | 37.933 * ± 3.193<br>p=9.134$\times 10^{-5}$ | 108.129 * ± 2.105<br>p=9.134$\times 10^{-5}$ |
| Comparison between standard DBS and DBS<sub>ca</sub> | | p=9.134$\times 10^{-5}$* | p=0.260 | p=9.134$\times 10^{-5}$* |

Furthermore, the strongest external input evoked significantly greater responses during DBS_*ca*_ compared to standard DBS (Fig. 5Bi, responses at 0.04 *µS* and Table 4B), suggesting that external inputs may be amplified by the presence of DBS_*ca*_ rather than masked. In the absence of an external input, the spike rates were suppressed by DBS_*ca*_ compared to standard DBS (Fig. 5Bi and Table 4B, responses at 0 *µS*), and this suppression due to DBS_*ca*_ might enable an increased signal-to-noise with the external input.

### 3.5 Biphasic pulse order may affect tremor reduction and response to external inputs

Lastly, we simulated biphasic DBS with the pulse order reversed from cathode-anode to anode-cathode (Fig. 6 and Table 5). While all previous simulations led with the cathodic phase (DBS_*ca*_), in Fig. 6 the anodic phase led. Tested at the same amplitudes as seen in Fig. 2, no significant differences were found between biphasic DBS types (Fig. 2Bi and Table 5Bi). In contrast, with intrinsic networks, DBS_*ac*_ led to slightly but significantly lower TFOs than DBS_*ca*_ from 1.00–2.00 mA (Fig. 6Bii and Table 5B). Comparison between extrinsic and intrinsic networks showed that DBS_*ac*_ reduced TFOs greater in the intrinsic network than the extrinsic network, as seen with standard DBS and DBS_*ca*_ (Table 5C).

**Figure 6:**
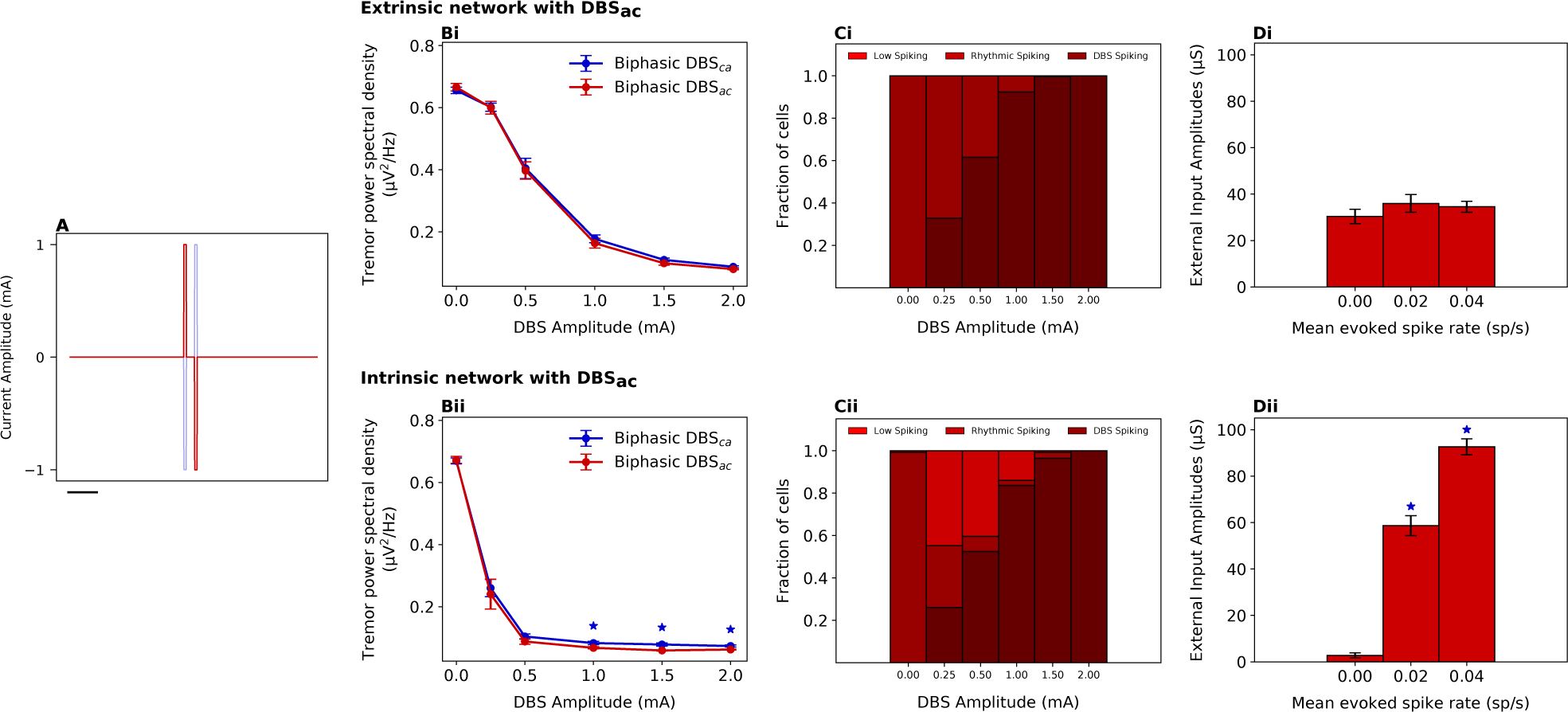
DBS_*ac*_ reduced tremor and affected input response during intrinsic TFOs. Pulse order of the biphasic stimulation was changed. Asterisks denote comparison to DBS_*ca*_. A. DBS_*ac*_ pulse shape (red) compared to DBS_*ca*_ shape (blue). Scale bar is 1 ms. B. Extrinsic network. Bi. DBS_*ac*_ reduced tremor similarly to DBS_*ca*_. Bii. Similar profile of spiking output during DBS_*ac*_ compared to standard DBS and DBS_*ca*_. Biii. No significant differences in evoked spike rate with external inputs during DBS_*ac*_ compared to DBS_*ca*_. C. Intrinsic network. Ci. DBS_*ac*_ reduced tremor significantly in (ii) intrinsic networks from 1.0– 2.0 mA. Cii. Among non-DBS cells, with significant reductions in overall tremor power at comparable levels, DBS_*ac*_ had fewer significantly suppressed cells than DBS_*ca*_. Ciii. At 0.02 *µS*, DBS_*ac*_ evoked lower rates than DBS_*ca*_, but at 0.04 *µS*, DBS_*ac*_ evoked lower rates than DBS_*ca*_.

**Table 5:**
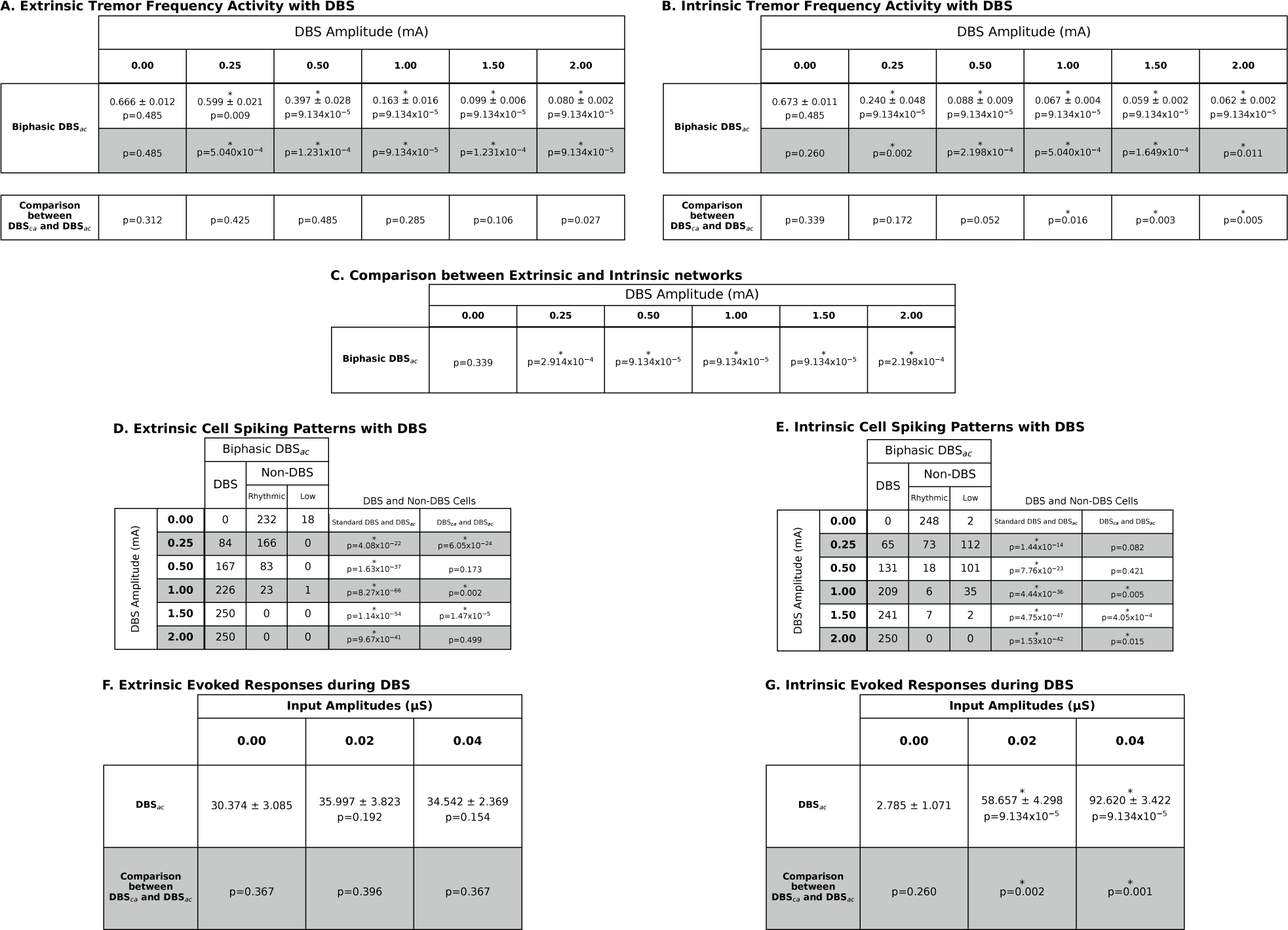
DBS_*ac*_ affected networks more. A. Extrinsic network. TFOs were significantly reduced with DBS_*ac*_ compared to baseline and were also significantly lower than standard DBS (gray row). DBS_*ac*_ did not change TFOs differently from DBS_*ca*_. B. Intrinsic network. TFOs were significantly reduced with DBS_*ac*_. Comparing to DBS_*ca*_, DBS_*ac*_ also significantly reduced TFOs from 1–2 mA. C. The intrinsic network demonstrated significantly lower TFOs compared to extrinsic. D–E. DBS_*ca*_ led to significantly more high frequency spiking cells compared to standard DBS. In the extrinsic network, DBS_*ac*_ generated significantly more high frequency spiking cells than DBS_*ca*_ at 0.25, 1.00, and 1.50 mA. In the intrinsic network, DBS_*ac*_ resulted in significantly more high frequency spiking cells than DBS_*ca*_ from 1.00–2.00 mA. F–G. At 0.02 *µS*, DBS_*ac*_ evoked significantly greater response than DBS_*ca*_, but at 0.04 *µS*, this was reversed, suggesting that phase order in biphasic DBS affects the ability for evoked inputs to be transmitted through VIM.

VIM cell spike outputs in the extrinsic network were also compared (Fig. 6C and Table 5D). The fraction of high frequency spiking cells was significantly greater for all tested amplitudes compared to standard DBS. This fraction was also significantly greater with DBS_*ac*_ than DBS_*ca*_, though only significantly at 0.25, 1.00, and 1.50 mA.

In the intrinsic network, DBS_*ac*_ led to significantly more high frequency spiking cells than standard DBS at all amplitudes and DBS_*ca*_ from 1.00–2.00 mA (Fig. 6Cii and Table 5E). For both intrinsic and extrinsic networks, these results suggest that DBS_*ac*_ may be better at driving high frequency activity in VIM cells than both standard DBS and DBS_*ca*_.

To determine whether non-tremor external input evoked spike rates were different during different biphasic DBS types, evoked rates were compared at each input level. For the extrinsic network, evoked rates during DBS_*ca*_ were not significantly different compared to DBS_*ac*_ (Fig. 6Di and Table 4, cf Fig. 5A). However, evoked responses in the intrinsic network were significantly different for DBS_*ac*_ compared to DBS_*ca*_. At 0.02 *µS*, the evoked responses during DBS_*ac*_ was significantly less than that during DBS_*ca*_ (Fig. 6Dii and Table 5G, cf Fig. 5Bi). But at 0.04 *µS*, evoked inputs during DBS_*ac*_ were significantly greater than those during DBS_*ca*_, demonstrating that the evoked response during different biphasic pulse sequences were nonlinear with respect to the synaptic input strength. These results raise the possibility that different pulse orders may be beneficial in distinct scenarios, depending on the strength of inputs to the VIM network.

Both biphasic DBS pulses showed significantly greater evoked responses to the synaptic input compared to the standard DBS. Overall this suggests that, for TFOs that are intrinsically generated, biphasic pulses can simultaneously reduce tremor and enable non-tremor inputs to be transmitted more effectively compared to standard DBS.

## 4 Discussion

We simulated the effects of DBS in a biophysically detailed network a thalamic network generating TFOs to understand local cellular mechanisms of tremor reduction. How DBS reduced TFOs in the model depended on the generating mechanism, and biphasic DBS was more effective at reducing TFOs compared to standard DBS. These results suggested that the specific stimulation pulse patterns may affect VIM cells in different ways and may be leveraged toward improved treatment. Furthermore, nontremor external inputs evoked greater responses during biphasic DBS compared to standard DBS, a potential mechanism for reducing DBS-induced side effects.

### 4.1 Relation to prior work on tremor suppression

Why DBS in VIM works to reduce tremor in patients with ET is not fully understood, and there may be multiple, possibly independent processes occurring. The prevailing theory for tremor cessation is simply the reduction of tremor frequency output of the motor cortex [10,33]. Computational modeling evidence has suggested that the reduction of tremor frequency output might be related to overwhelming the thalamic circuit with high frequency axonal output [33] that may also lead to synaptic failure [42].

The optimal site of DBS stimulation for treatment of ET may be VIM [18] or potentially the cerebellothalamic tract entering the thalamus [44]. If true, this would suggest that the input from the cerebellum to the thalamus is giving rise to or at the very least promoting tremorgenic transmission, but it is still unclear if cerebellum is the principal driver of thalamic tremor activity and to what extent intrinsic mechanisms are involved [27]. Here, we explicitly modeled extrinsic and intrinsic TFOs using the same VIM-TRN network, with reciprocal excitatory and inhibitory connections. The network architecture and responses to inputs provided a crucial framework to understanding how different mechanisms specifically generating tremor frequency activity would be perturbed by different types of DBS.

In the model, biphasic pulse shapes were more effective at reducing the TFOs in the spiking output of VIM compared to standard DBS (Fig. 2, Fig. 6B). A closer look at the individual VIM cell activities suggested that this might be due to the overall number of cells participating in “pathological” oscillations, which was far greater in standard DBS.

In these simulations, DBS_*ac*_ reduced tremor slightly more effectively than other pulse patterns, principally by driving more cells to high spike rates. Yet standard DBS reduced tremor activity by not only increasing high spike rate activity but also by decorrelating rhythmic activity within VIM, while biphasic DBS led to a combination of both. DBS also suppressed output in certain VIM cells, which involved T-type Ca^2+^ inactivation (Fig. 3–4). These results suggest that a “sweet spot” may exist in DBS stimulation, in which the number of rhythmic cells is reduced to control tremor, while a substantial population of cells shift from rhythmicity to quiescence via T-type Ca^2+^ inactivation. These dynamics were principally manifested by biphasic DBS and suggest a possible strategy of refining thalamic modulation techniques to suppress rhythmic bursting activity instead of entraining high frequency spiking output.

Here, we focused on identifying whether biphasic pulse patterns could differentially affect the biophysics of individual cells in the VIM network to reduce tremor. We did not test all possible parameters related to DBS. Prior experimental and computational work has shown that the frequency [38], periodicity [9,48], pulse width [23], and varied pulse shapes [16,20] all may affect the efficacy of DBS stimulation. Our results suggest the biophysical mechanisms of generation of TFOs have a direct influence on the efficacy of DBS and that such patterns should be reexamined in the context of our biophysical model. If biphasic pulses still reduce tremor effectively at lower frequencies than standard DBS, while suppressing spiking in VIM cells, this would represent another avenue of improvement for DBS efficiency.

In particular, we conjecture that pulse width may lead to increased excitability of VIM cells when coupled to the biphasic pulse sequences explored here. The T-type Ca^2+^ channels in these cells may be activated when longer pulses are able to hold VIM cell membrane potentials at levels that might amplify the gating variables necessary to cause strong T-type Ca^2+^ currents, similar to the mechanism observed in TRN cells seen in Fig. 3 in [27]. Longer pulse widths would therefore lead to entrainment at DBS frequencies, resulting in reduced tremor activity.

In all of our simulations, we fixed the recording electrode locations at 250 *µm* away from the center of mass of the VIM, but it is well known that the stimulation strength is highly dependent on the geometry of the cells with respect to the electrodes [33]. We varied the strength of the stimulation at this location, which suggests that some of the effects reported—especially the recruitment of different types of spiking in VIM cells with different stimulation patterns—would also hold true for different stimulation locations, at least along the y-axis of our network (see Fig. 1C–D).

### 4.2 Focal modulation of membrane biophysics with electrical stimulation

Prior computational modeling has implicated the role of T-type Ca^2+^ channels in movement disorder pathophysiology [43]. Our standard DBS model elicited both high spike rates as well as uncorrelated but rhythmic spiking in cells that did not follow the DBS stimulation (Fig. 2). These results taken together suggest that reducing or leveraging the biophysics of T-type Ca^2+^ currents, particularly as they relate to tremor frequency oscillations, may be crucial in reducing pathological tremor activity from being manifested in the VIM, with potential consequences for physiological tremor in patients. Medications targeting T-type Ca^2+^ channel pathology are likely to be broadly acting. In contrast, the effect of DBS_*ca*_ and DBS_*ac*_ pulses in the present work demonstrated that manipulating T-type Ca^2+^ channels may not only reduce tremor frequency activity that ultimately reduces tremor in ET but also does so in a focal, non-pharmacological way, providing a targeted therapy whose activation volume is limited to that of the stimulation.

### 4.3 Implications of biphasic DBS on effective throughput of relevant cerebellar signals

The cerebellum is thought to contribute to motor planning and adaptation, and disruption of the cerebellothalamocortical pathway via lesion or DBS has been shown to impair these functions [10]. The role of cerebellar activity raises the question of whether the presence of tremor frequency oscillations in the VIM affect normal non-tremor inputs from the cerebellum that might mediate motor adaptation. The impairment of motor adaptation by standard DBS also raises the question of whether it is possible to design DBS stimulation that simultaneously reduces tremor and permits these inputs to be expressed. Our results suggest that biphasic DBS is one such pattern that may alleviate tremor while allowing relevant cerebellar signals to be propagated through thalamus to cortex.

In this study, the tremor frequency generating network was assumed to be the same that received and processed non-tremor cerebellar inputs, yet it is not fully known whether these networks serve the same functions in the VIM. If these networks are not the same, then the question would shift to whether ongoing DBS pulse sequences elicit activity in non-tremor cells similar to that observed here. In this scenario, electrode placement or patterns of stimulation that optimally targeted non-tremor cells might act to help cerebellar inputs to be transmitted, but ongoing tremor might still be expressed through a separate channel through the VIM. Only the biphasic stimulation simulated here addressed both the tremor and non-tremor components without condition, and while tremor was reduced in both the extrinsic and intrinsic networks, inputs were only affected in the intrinsic network. Our results suggest that it is possible that these DBS pulses also silence neighboring networks of non-tremor cells, also enabling these inputs to be represented.

The phase order of biphasic DBS pulses also affected the amplitude of the external, non-tremor cerebellar input into the intrinsic network. While the 0.02 *µS* input created a significantly greater evoked response during DBS_*ac*_, the stronger input created the opposite effect (cf. Fig. 6Dii and Fig. 5Bi). One possible utility of this result could be a DBS design with rapid switching of phase order based on the patient’s needs, for instance with DBS_*ac*_ at low levels to suppress tremor efficiently and a switch to DBS_*ca*_ when the patient is speaking or performing fine motor tasks.

Similar approaches to understanding the effect of DBS on the biophysics of networks may also help to inform mechanisms of action for DBS in other diseases. DBS has been deployed at various targets for treatment of a host of diseases including Parkinson’s disease [15], Alzheimer’s disease [31,40], obsessive-compulsive disorder [17,35], and depression [21], and the treatment is being used for other indications as well. In Parkinson’s disease, DBS in the subthalamic nucleus (STN) reduces symptoms like tremor but may lead to cognitive side effects [36]. Prior computational modeling work has suggested that DBS in the subthalamic nucleus tonically activated the globus pallidus internus (GPi), which led to tonic inhibition of the pallidal-receiving thalamus, suggested to be favorable to phasic inhibition that led to pathological oscillatory activity [43]. If the present model results also applied to the T-type Ca^2+^ channels found in STN [49], where biphasic DBS led to both DBS-following cells and spike-suppressed cells, the DBS-following cells may still lead to tonic inhibition from GPi but also allow other inputs into STN to be more faithfully represented. Preliminary evidence has demonstrated efficacy of biphasic DBS in STN for the treatment of PD [3], but further work is necessary to demonstrate whether these particular mechanisms are involved.

## Conclusions

DBS has been an effective though imperfect therapy, and its future improvement relies on a more thorough understanding of its targets and the mechanisms of stimulation. Our model results demonstrated that the mechanisms of DBS to reduce TFOs were dependent upon how the TFOs were generated biophysically. Biphasic DBS was more effective in reducing TFOs in the model and may be leveraging T-type Ca^2+^ dynamics to suppress activity in VIM, facilitating the transmission of non-tremor activity from cerebellum to cortex. Together, the model results suggested modulation of specific biophysical mechanisms toward designing improved treatments for ET and other diseases.

## 5 Acknowledgements

This work was funded in parts by NIMH 5T32MH019118-23 (supporting SL), NIMH R01 MH106174 (SRJ), Brown University BioMed Dean’s Emerging Areas of New Science Award (SRJ, WFA), the Brown Institute for Brain Sciences and Norman Princes Neuroscience Institute New Frontiers Fund (SRJ, WFA), NIH COBRE P20 GM103645 (WFA), and NIH R01 MH115035 (WFA).

